## Supplemental Document for "Common synaptic phenotypes arising from diverse mutations in the human NMDA receptor subunit GluN2A"

### Supplemental Information

#### Figures S1.1 – S1.11: Summary of Linear Mixed Models (LMM)

The top of each of the following figures includes: the `lmer` (Wilkinson-Rogers-Pinheiro-Bates) model formula, an ANOVA table with $F(df, df_{res})$ statistics, *p*-values and Bayes factors for hypothesis tests of interests on fixed effects, and a table of group sample size and intraclass correlation (ICC) for nuisance random effects (and the residual, unexplained error). The bottom of each of the following figures (1.1-1.11) includes a series of plots for assessing the validity of assumptions made by the LMM: **a**) Standardized residuals plotted against the fitted values with smoothed conditional mean (red) and conditional median accompanied by lower and upper quartiles (blue) ; **b**) Histogram of residual overlaid with fitted Gaussian distribution (red) and kernel density estimate (blue); **c**) quantile-quantile (Q-Q) plot with 95% confidence bands; and **d**) Stem-and-leaf plot of Cook's distance.

##### *Figure S1.1. NMDA-EPSC peak amplitudes in GluN2A/B double KO neurons rescued with GluN2A mutants*

***Top***. Table summary of a Linear Mixed Model (LMM) predicting NMDA-EPSC peak amplitude from the co-transfection of *GRIN2A* mutants and Cre-GFP into CA1 neurons of *grin2a*$^{fl/fl}$*2b*$^{fl/fl}$ neurons. Fixed effects are summarised as an ANOVA table with the interaction term decomposed into orthogonal contrasts (A-E, see Table S1). The *p*-value and Bayes factor for the interaction term omnibus test indicate a highly significant effect of transfecting GluN2A mutants on the peak amplitude of NMDA-EPSCs in GluN2A/B double knockout (KO) neurons. Of the variance not accounted for by the fixed effects, 41% and 59% was explained by variability between recording pairs or residual (unexplained) variance respectively. ***Bottom***. Model residuals appeared to be homoscedastic (**a**) and normally distributed (**b**, **c**), without outliers (**a, c**) or overly influential data points (**d**). A breakdown of samples sizes is reported in Table S4.

| **Response** | | | | | |
|---|---|---|---|---|---|
| NMDA-EPSC peak amplitude (peak) | | | | | |
| **Formula** | | | | | |
| $\log(\boldsymbol{peak}) \sim mutation * transfection + (1 \vert animal/slice/pair)$ | | | | | |
| **Source (Fixed)** | ***F*** | ***df*** | ***df$_{res}$*** | ***p*** | ***BF$_{10}$*** |
| mutation | 7.89 | 5 | 8.8 | | |
| transfection | 433.89 | 1 | 87 | | |
| mutation:transfection | 11.35 | 5 | 87 | <.001 | 4.14E+06 |
| A. WT *vs.* K669N, L812M, C436R, T531M, R518H | 31.22 | 1 | 87 | <.001 | |
| B. K669N, L812M *vs.* C436R, T531M, R518H | 25.94 | 1 | 87 | <.001 | |
| C. R518H *vs.* C436R, T531M | 0.01 | 1 | 87 | .93 | |
| D. C436R *vs.* T531M | 0.14 | 1 | 87 | .71 | |
| E. K669N *vs.* L812M | 0.00 | 1 | 87 | .98 | |
| **Source (random)** | ***N*** | **ICC** | | | |
| pair:(slice:animal) | 93 | .41 | | | |
| slice:animal | 54 | .00 | | | |
| animal | 16 | .00 | | | |
| Residual | | .59 | | | |

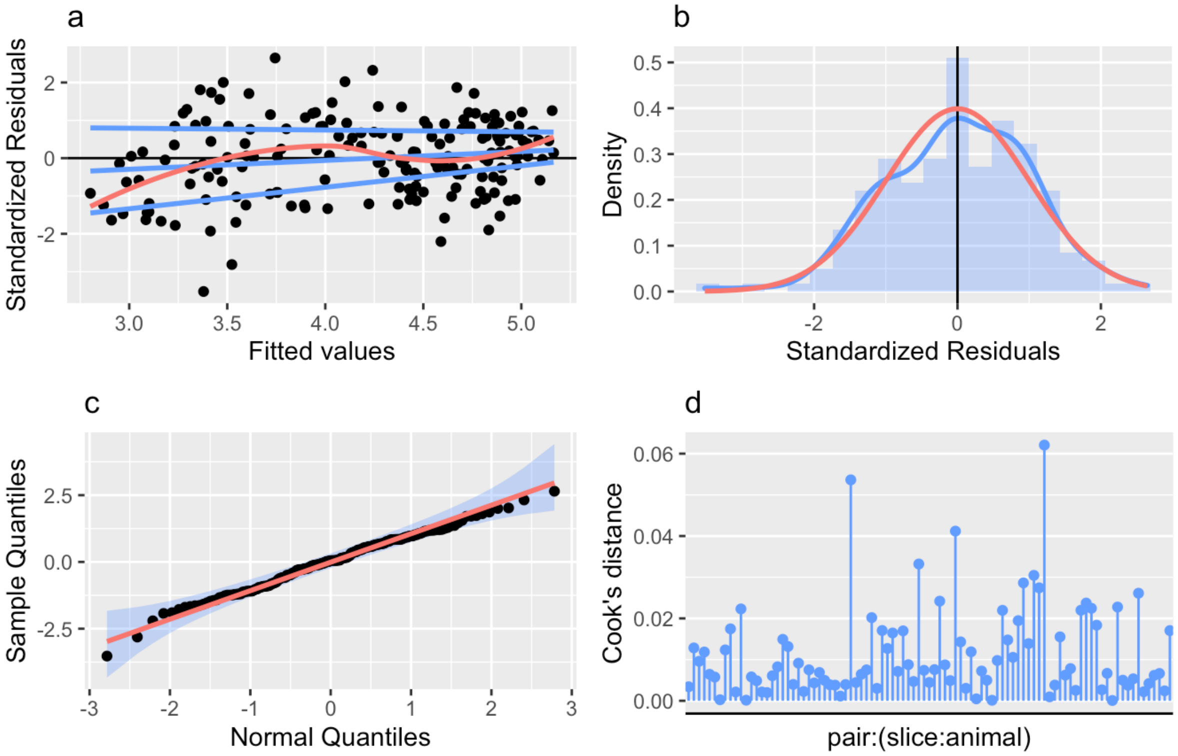

***Figure S1.2. NMDA-EPSC decays in GluN2A/B double KO neurons rescued with GluN2A GOF mutants***

***Top***. Table summary of a LMM predicting NMDA-EPSC decay time constant from the co-transfection of *GRIN2A* mutants and Cre-GFP in CA1 neurons of *grin2a$^{fl/fl}$2b$^{fl/fl}$* neurons. Fixed effects are summarised as an ANOVA table with the interaction followed up with posthoc pairwise comparisons. The *p*-value and Bayes factor for the interaction term omnibus test indicate a highly significant effect of transfecting GluN2A mutants on the decay time constant of NMDA-EPSCs in GluN2A/B double knockout (KO) neurons. Of the variance not accounted for by the fixed effects, 20%, 7%, 27% and 46% was explained by variability between recording pairs, slices, animals and by residual (unexplained) variance respectively. ***Bottom***. Model residuals appeared to be homoscedastic (**a**) and normally distributed (**b**, **c**), without overt outliers (**a**,**c**) or overly influential data points (**d**). A breakdown of samples sizes is reported in Table S4.

| **Response** | | | | | |
|---|---|---|---|---|---|
| NMDA-EPSC decay time constant (decay) | | | | | |
| **Formula** | | | | | |
| $\log(\boldsymbol{decay}) \sim mutation * transfection + (1\|animal/slice/pair)$ | | | | | |
| **Source (Fixed)** | ***F*** | ***df*** | ***df$_{res}$*** | ***p*** | ***BF$_{10}$*** |
| mutation | 69.01 | 2 | 5.71 | | |
| transfection | 19.06 | 1 | 49 | | |
| mutation:transfection | 190.3 | 2 | 49 | <.001 | 7.61E+40 |
| **Posthoc test** | ***t*** | | ***df*** | ***p$_{adj}$*** | |
| WT *vs.* K669N | 5.3 | | 49 | <.001 | |
| WT *vs.* L812M | 13.29 | | 49 | <.001 | |
| K669N *vs.* L812M | 9.1 | | 49 | <.001 | |
| **Source (random)** | ***N*** | **ICC** | | | |
| pair:(slice:animal) | 52 | .20 | | | |
| slice:animal | 29 | .07 | | | |
| animal | 9 | .27 | | | |
| Residual | | .46 | | | |

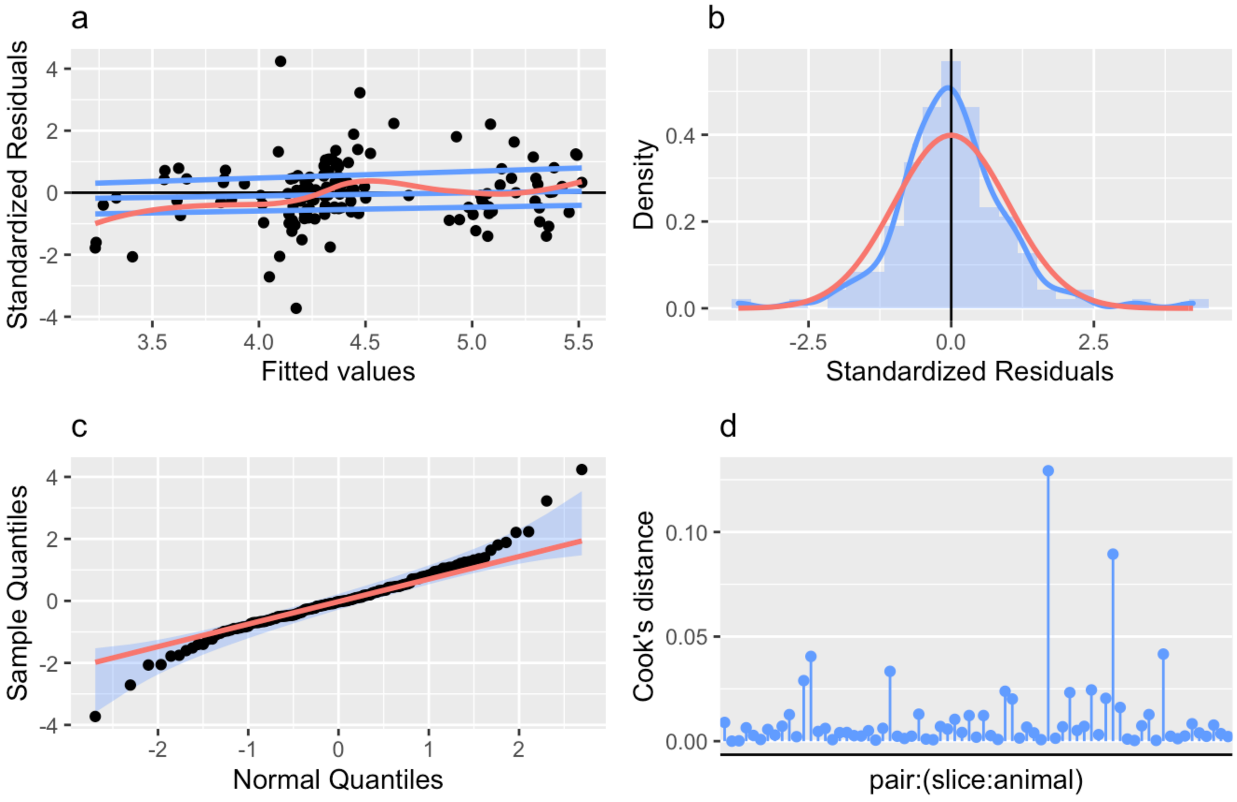

***Figure S1.3. Synaptic GFP-GluN1 expression rescued by expression of GluN2A mutants.***

***Top***. Table summary of a LMM predicting fluorescence intensity from GFP-GluN1 and Homer1c expressed in cultured hippocampal neurons co-transfected with different *GRIN2A* mutants. Fixed effects are summarised as an ANOVA table with the interaction term decomposed into orthogonal contrasts (A-E, see Table S1). The *p*-value and Bayes factor for the interaction term omnibus test indicate a highly significant effect of GluN2A mutants on the fluorescence of GFP-GluN1 relative to Homer1c-tdTomato. Of the variance not accounted for by the fixed effects, 19, 9 and 72% was explained by variability between regions of interest (roi), experiments or residual (unexplained) variance respectively. ***Bottom***. Model residuals appeared to be mostly homoscedastic (**a**) and normally distributed (**b**, **c**), without overt outliers (**a, c**) or overly influential data points (**d**). A breakdown of samples sizes is reported in Table S4.

| **Response** | | | | | |
|---|---|---|---|---|---|
| Fluorescence intensity (fluorescence) | | | | | |
| **Formula** | | | | | |
| $\log(\boldsymbol{fluorescence}) \sim mutation * protein + (1 \vert expt/roi)$ | | | | | |
| **Source (Fixed)** | ***F*** | ***df*** | $\boldsymbol{df_{res}}$ | ***p*** | $\boldsymbol{BF_{10}}$ |
| mutation | 21.02 | 5 | 126 | | |
| protein | 1902 | 1 | 185 | | |
| mutation:protein | 10.29 | 5 | 185 | <.001 | 4.82E+04 |
| A. WT *vs*. K669N, L812M, C436R, T531M, R518H | 12.71 | 1 | 185 | <.001 | |
| B. K669N, L812M *vs.* C436R, T531M, R518H | 6.9 | 1 | 185 | .009 | |
| C. R518H *vs.* C436R, T531M | 21.65 | 1 | 185 | <.001 | |
| D. C436R *vs.* T531M | 7.45 | 1 | 185 | .007 | |
| E. K669N *vs.* L812M | 9.32 | 1 | 185 | .003 | |
| **Source (random)** | ***N*** | **ICC** | | | |
| roi:expt | 191 | .19 | | | |
| expt | 8 | .09 | | | |
| Residual | | .72 | | | |

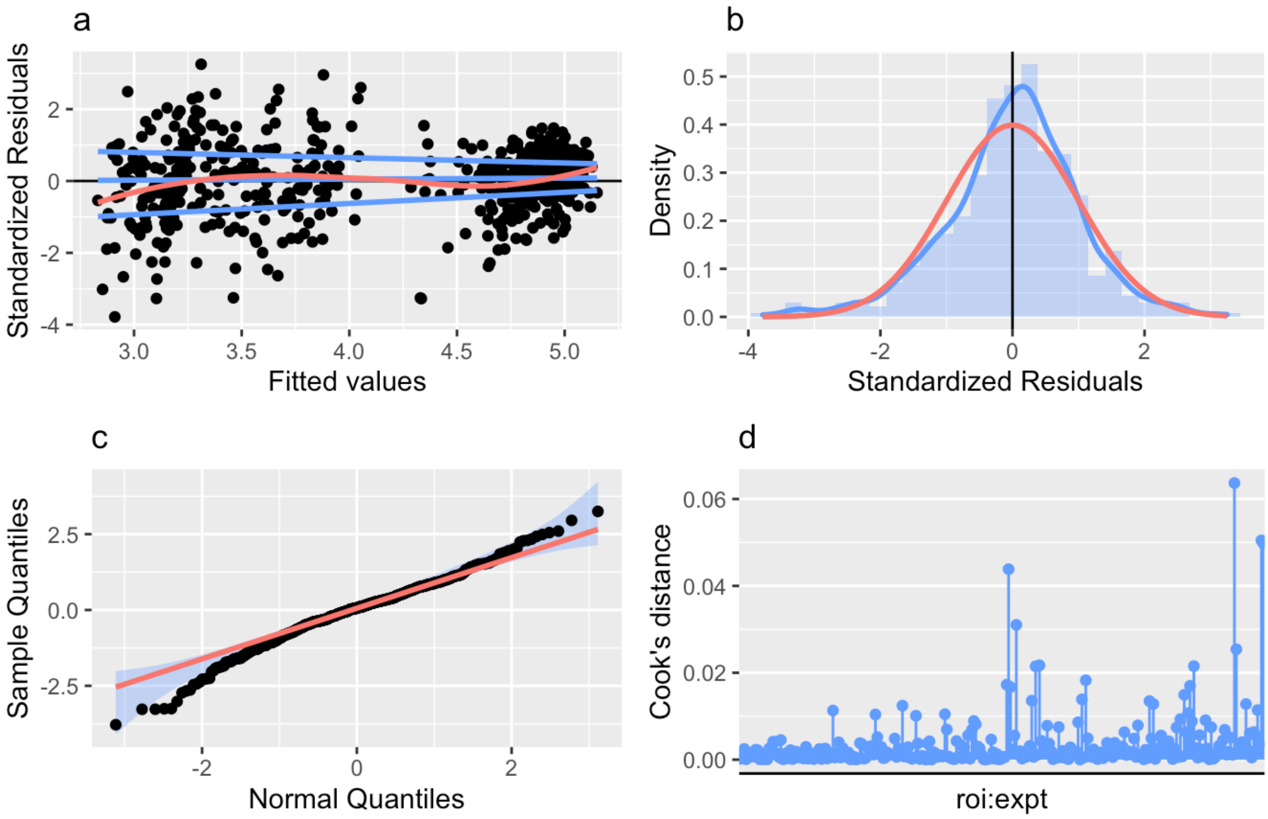

***Figure S1.4. NMDA-EPSC peak amplitudes in GluN2A knockout neurons rescued with GluN2A mutants.***

***Top***. Table summary of a LMM predicting NMDA-EPSC peak amplitude from the co-transfection of *GRIN2A* mutants and Cre-GFP in CA1 neurons of *grin2a*$^{fl/fl}$ neurons. Fixed effects are summarised as an ANOVA table with the interaction term decomposed into orthogonal contrasts (A-E, see Table S1). The Bayes factor for the interaction term omnibus test provides evidence more in favour of no effect of GluN2A mutants on the peak amplitude of NMDA-EPSCs in GluN2A knockout (KO) neurons. Of the variance not accounted for by the fixed effects, 24%, 22%, 37% and 16% was explained by variability between recording pairs, slices, animals and residual (unexplained) variance respectively. ***Bottom***. Model residuals appeared to be homoscedastic (**a**) and normally distributed (**b**, **c**), without outliers (**a**,**c**) or influential data points (**d**). A breakdown of samples sizes is reported in Table S4.

| **Response** | | | | | |
|---|---|---|---|---|---|
| NMDA-EPSC peak amplitude (peak) | | | | | |
| **Formula** | | | | | |
| $\log(\boldsymbol{peak}) \sim mutation * transfection + (1\vert animal/slice/pair)$ | | | | | |
| **Source (Fixed)** | ***F*** | ***df*** | ***df$_{res}$*** | ***p*** | ***BF$_{10}$*** |
| mutation | 0.72 | 5 | 19.8 | | |
| transfection | 78.02 | 1 | 119 | | |
| mutation:transfection | 0.87 | 5 | 119 | .50 | 0.0838 |
| A. WT vs K669N, L812M, C436R, T531M, R518H | 2.47 | 1 | 119 | .12 | |
| B. K669N, L812M vs. C436R, T531M, R518H | 1.88 | 1 | 119 | .17 | |
| C. R518H vs. C436R, T531M | 0.25 | 1 | 119 | .62 | |
| D. C436R vs. T531M | 0.00 | 1 | 119 | .99 | |
| E. K669N vs. L812M | 0.14 | 1 | 119 | .71 | |
| **Source (random)** | ***N*** | **ICC** | | | |
| pair:(slice:animal) | 125 | .24 | | | |
| slice:animal | 78 | .22 | | | |
| animal | 27 | .37 | | | |
| Residual | | .16 | | | |

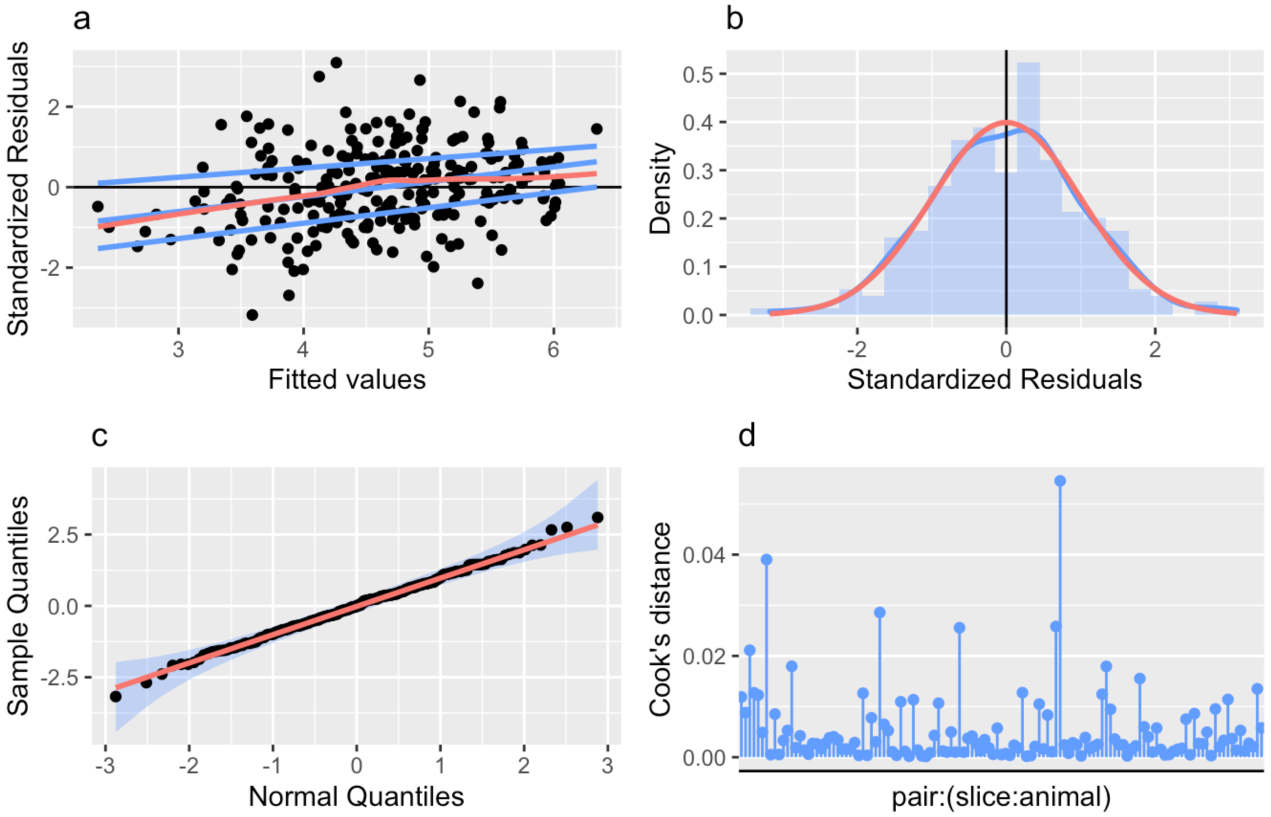

##### *Figure S1.5. NMDA-EPSC decays in GluN2A knockout neurons rescued with GluN2A mutants.*

***Top***. Table summary of a LMM predicting NMDA-EPSC decay time constant from the co-transfection of *GRIN2A* mutants and Cre-GFP in CA1 neurons of *grin2a*$^{fl/fl}$ neurons. Fixed effects are summarised as an ANOVA table with the interaction term decomposed into orthogonal contrasts (A-E, see Table S1). The *p*-value and Bayes factor for the interaction term omnibus test indicate a highly significant effect of transfecting GluN2A mutants on the decay time constant of NMDA-EPSCs in GluN2A knockout (KO) neurons. Of the variance not accounted for by the fixed effects, 4%, 12%, 16% and 68% was explained by variability between recording pairs, slices, animals and residual (unexplained) variance respectively. ***Bottom***. Model residuals appeared to be homoscedastic (**a**) and normally distributed (**b**, **c**), without outliers (**a**,**c**) or influential data points (**d**). A breakdown of samples sizes is reported in Table S4.

**Response**

NMDA-EPSC decay time constant (decay)

**Formula**

$\log(\boldsymbol{decay}) \sim mutation * transfection + (1|animal/slice/pair)$

| **Source (Fixed)** | ***F*** | ***df*** | ***df$_{res}$*** | ***p*** | ***BF$_{10}$*** |
|---|---|---|---|---|---|
| mutation | 3.06 | 5 | 19.3 | | |
| transfection | 179.6 | 1 | 119 | | |
| mutation:transfection | 12.09 | 5 | 119 | <.001 | 3.43E+08 |
| A. WT vs K669N, L812M, C436R, T531M, R518H | 37.09 | 1 | 119 | <.001 | |
| B. K669N, L812M vs. C436R, T531M, R518H | 28.34 | 1 | 119 | <.001 | |
| C. R518H vs. C436R, T531M | 0.13 | 1 | 119 | .72 | |
| D. C436R vs. T531M | 0.23 | 1 | 119 | .63 | |
| E. K669N vs. L812M | 0.12 | 1 | 119 | .73 | |
| **Source (random)** | ***N*** | **ICC** | | | |
| pair:(slice:animal) | 125 | .04 | | | |
| slice:animal | 78 | .12 | | | |
| animal | 27 | .16 | | | |
| Residual | | .68 | | | |

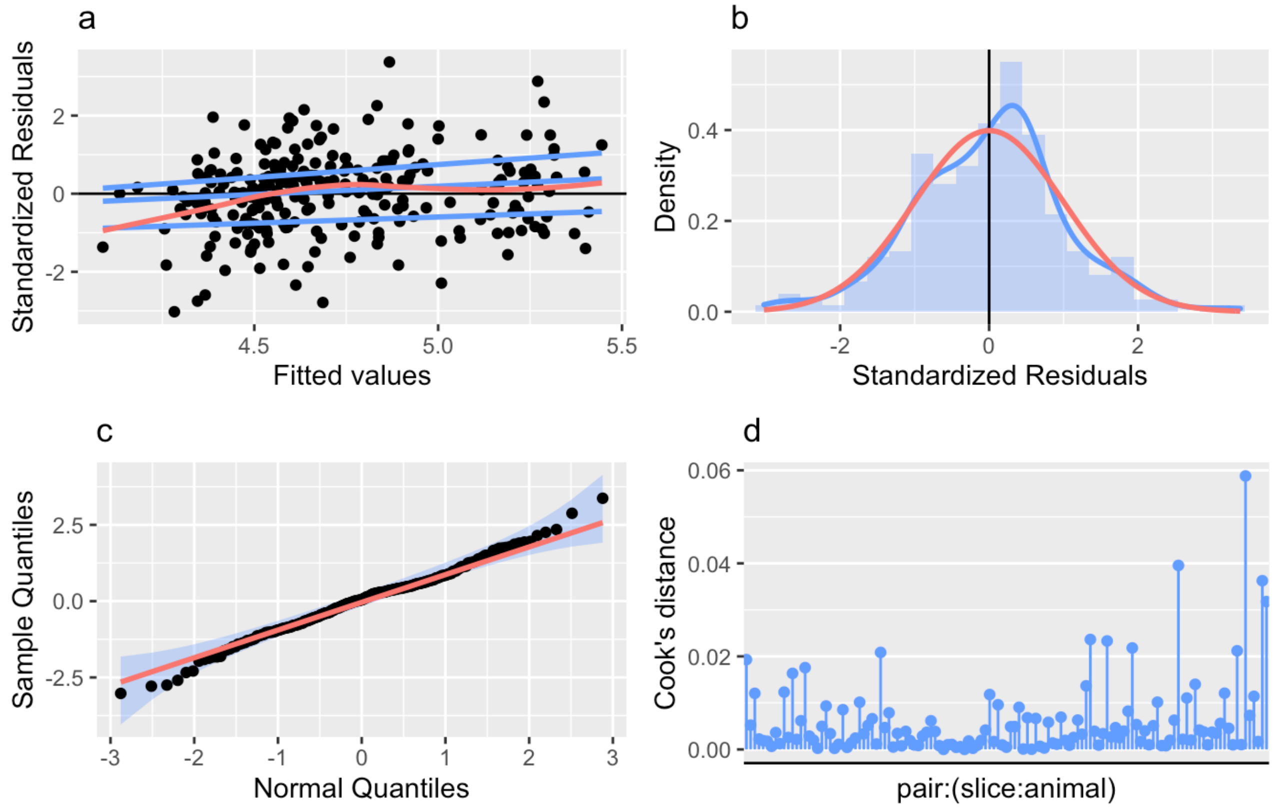

***Figure S1.6. NMDA-EPSC charge transfer in GluN2A knockout neurons rescued with GluN2A mutants.***

***Top***. Table summary of a LMM predicting NMDA-EPSC charge transfer from the co-transfection of *GRIN2A* mutants and Cre-GFP in CA1 neurons of *grin2a*$^{fl/fl}$ neurons. Fixed effects are summarised as an ANOVA table with the interaction term decomposed into orthogonal contrasts (A-E, see Table S1). The Bayes factor for the interaction term omnibus test provides weak evidence for no effect of GluN2A mutants on the charge transfer of NMDA-EPSCs in GluN2A knockout (KO) neurons. Of the variance not accounted for by the fixed effects, 33%, 22%, 27% and 18% was explained by variability between recording pairs, slices, animals and residual (unexplained) variance respectively. ***Bottom***. Model residuals appeared to be homoscedastic (**a**) and normally distributed (**b**, **c**), without outliers (**a, c**) or influential data points (**d**). A breakdown of samples sizes is reported in Table S4.

| **Response** | | | | | |
|---|---|---|---|---|---|
| NMDA-EPSC charge transfer (charge) | | | | | |
| **Formula** | | | | | |
| $\log(\boldsymbol{charge}) \sim mutation * transfection + (1 \vert animal/slice/pair)$ | | | | | |
| **Source (Fixed)** | ***F*** | ***df*** | ***df$_{res}$*** | ***p*** | ***BF$_{10}$*** |
| mutation | 1.05 | 5 | 19.5 | | |
| transfection | 0 | 1 | 119 | | |
| mutation:transfection | 2.13 | 5 | 119 | .07 | 0.632 |
| A. WT vs K669N, L812M, C436R, T531M, R518H | 4.18 | 1 | 119 | .04 | |
| B. K669N, L812M vs. C436R, T531M, R518H | 4.26 | 1 | 119 | .04 | |
| C. R518H vs. C436R, T531M | 2.47 | 1 | 119 | .12 | |
| D. C436R vs. T531M | 0.17 | 1 | 119 | .68 | |
| E. K669N vs. L812M | 0.32 | 1 | 119 | .57 | |
| **Source (random)** | ***N*** | **ICC** | | | |
| pair:(slice:animal) | 125 | .33 | | | |
| slice:animal | 78 | .22 | | | |
| animal | 27 | .27 | | | |
| Residual | | .18 | | | |

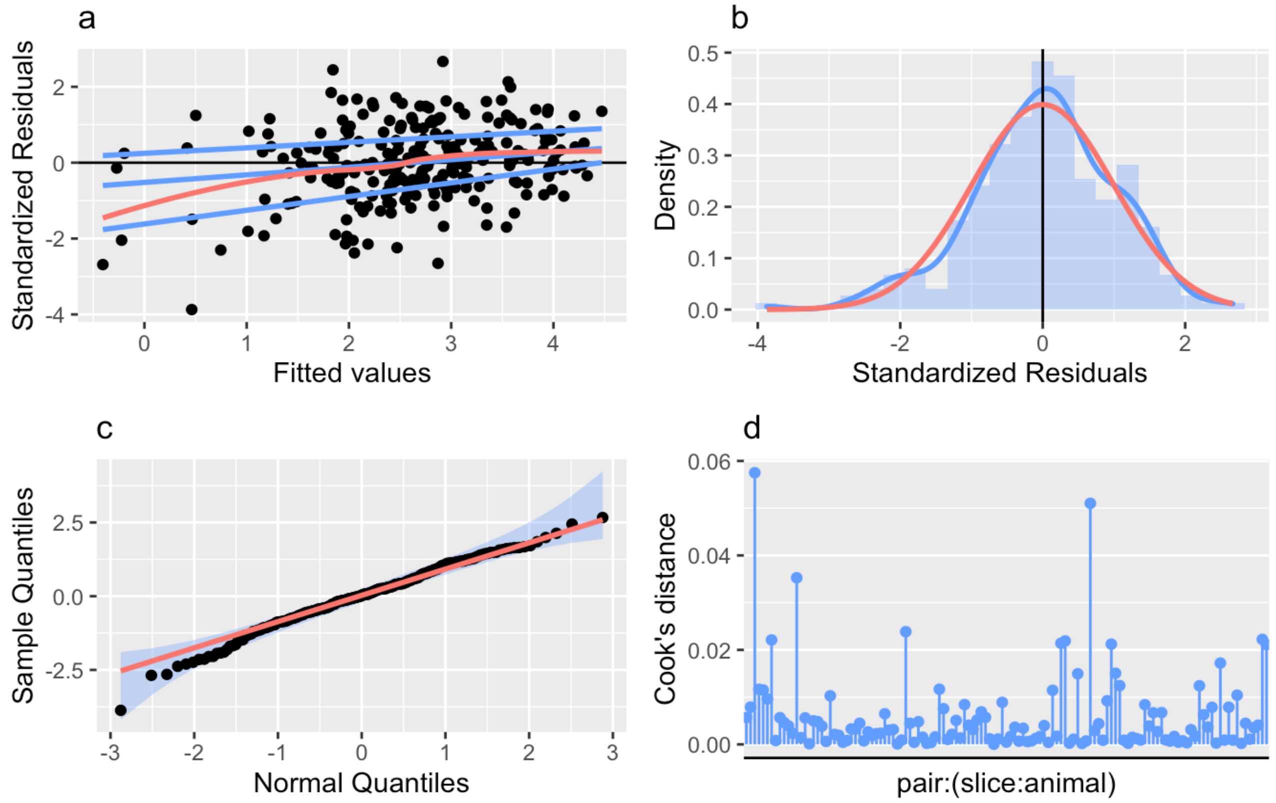

##### *Figure S1.7. AMPA-EPSC peak amplitudes in GluN2A knockout neurons rescued with GluN2A mutants.*

***Top***. Table summary of a LMM predicting AMPA-EPSC peak amplitude from the co-transfection of *GRIN2A* mutants and Cre-GFP in CA1 neurons of *grin2a*$^{fl/fl}$ neurons. Fixed effects are summarised as an ANOVA table with the interaction term decomposed into orthogonal contrasts (A-E, see Table S1). The Bayes factor for the interaction term omnibus test provides reasonable evidence for no effect of GluN2A mutants on the peak amplitude of AMPA-EPSCs in GluN2A knockout (KO) neurons. Of the variance not accounted for by the fixed effects, 27%, 40% and 33% was explained by variability between recording pairs, animals and residual (unexplained) variance respectively. ***Bottom***. Model residuals appeared to be homoscedastic (**a**) and normally distributed (**b**, **c**), without outliers (**a, c**) or influential data points (**d**). A breakdown of samples sizes is reported in Table S4.

| **Response** | | | | | |
|---|---|---|---|---|---|
| AMPA-EPSC peak amplitude (peak) | | | | | |
| **Formula** | | | | | |
| $\log(\boldsymbol{peak}) \sim mutation * transfection + (1\|animal/slice/pair)$ | | | | | |
| **Source (Fixed)** | ***F*** | ***df*** | ***df$_{res}$*** | ***p*** | ***BF$_{10}$*** |
| mutation | 2.66 | 5 | 19.95 | | |
| transfection | 2.38 | 1 | 119 | | |
| mutation:transfection | 1.49 | 5 | 119 | .20 | 0.267 |
| A. WT vs K669N, L812M, C436R, T531M, R518H | 0.04 | 1 | 119 | .84 | |
| B. K669N, L812M vs. C436R, T531M, R518H | 7.23 | 1 | 119 | .008 | |
| C. R518H vs. C436R, T531M | 0.15 | 1 | 119 | .70 | |
| D. C436R vs. T531M | 0.05 | 1 | 119 | .82 | |
| E. K669N vs. L812M | 0.01 | 1 | 119 | .93 | |
| **Source (random)** | ***N*** | **ICC** | | | |
| pair:(slice:animal) | 125 | .27 | | | |
| slice:animal | 78 | .00 | | | |
| animal | 27 | .40 | | | |
| Residual | | .33 | | | |

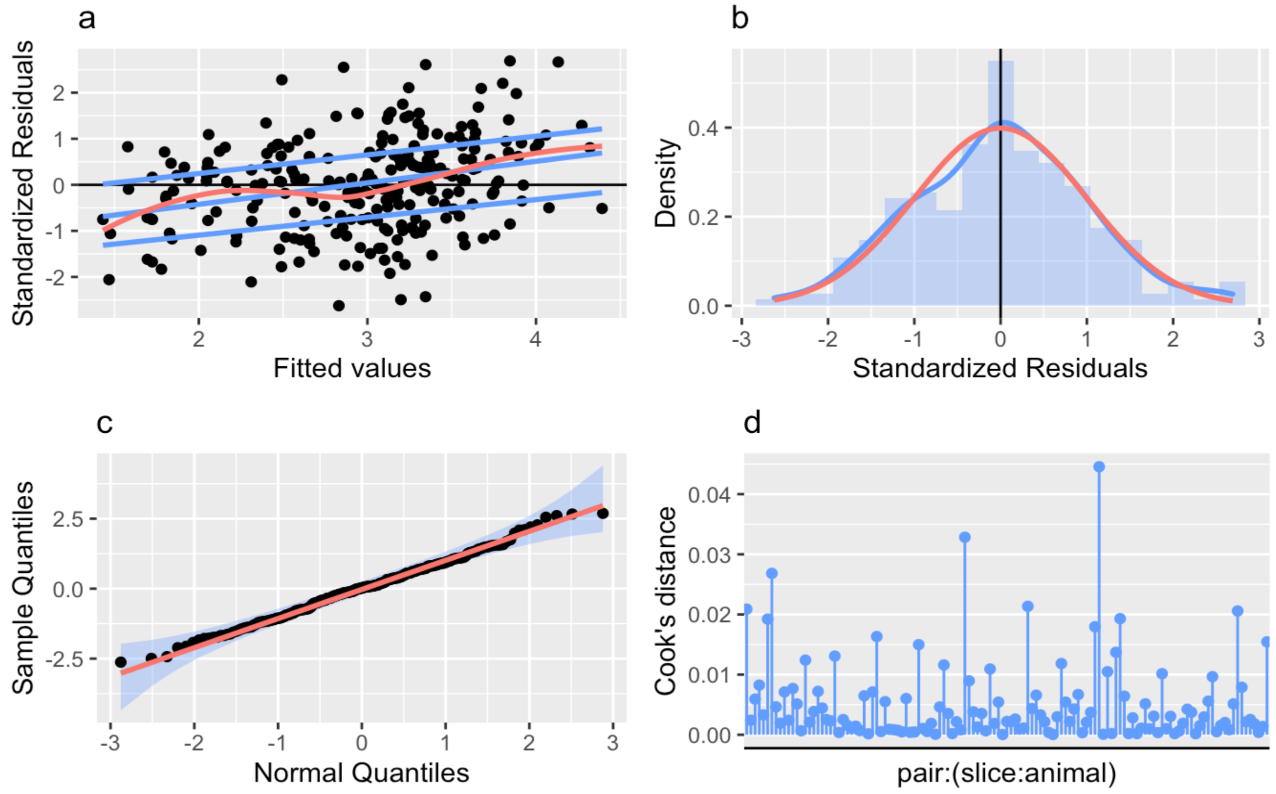

***Figure S1.8. NMDA-EPSC charge transfer in neurons expressing GluN2A knockout alleles.***

***Top***. Table summary of a LMM predicting NMDA-EPSC charge transfer from the transfection of Cre-GFP in CA1 neurons of *grin2a*$^{fl/fl}$ neurons. Fixed effects are summarised as an ANOVA table with the interaction term split into polynomial contrasts and followed up with posthoc pairwise comparisons. The Bayes factor for the interaction term omnibus test provides reasonable evidence for no effect of loss-of-function (null) GluN2A alleles on the charge transfer of NMDA-EPSCs. Of the variance not accounted for by the fixed effects, 1%, 32%, 31% and 36% was explained by variability between recording pairs, slices, animals and residual (unexplained) variance respectively. ***Bottom***. Model residuals appeared to be homoscedastic (**a**) and normally distributed (**b**, **c**), without outliers (**a, c**) or influential data points (**d**). A breakdown of samples sizes is reported in Table S4.

| **Response** | | | | | |
|---|---|---|---|---|---|
| NMDA-EPSC charge transfer (charge) | | | | | |
| **Formula** | | | | | |
| $\log(\boldsymbol{charge}) \sim genotype * transfection + (1|animal/slice/pair)$ | | | | |
| **Source (Fixed)** | ***F*** | ***df*** | ***df$_{res}$*** | ***p*** | ***BF$_{10}$*** |
| genotype | 0.09 | 2 | 10.09 | | |
| transfection | 0.01 | 1 | 96 | | |
| genotype:transfection | 1.21 | 2 | 96 | .30 | 0.277 |
| Linear | 2.39 | 1 | 96 | .13 | |
| Quadratic | 0.01 | 1 | 96 | .91 | |
| **Source (random)** | ***N*** | **ICC** | | | |
| pair:(slice:animal) | 125 | .01 | | | |
| slice:animal | 78 | .32 | | | |
| animal | 27 | .31 | | | |
| Residual | | .36 | | | |

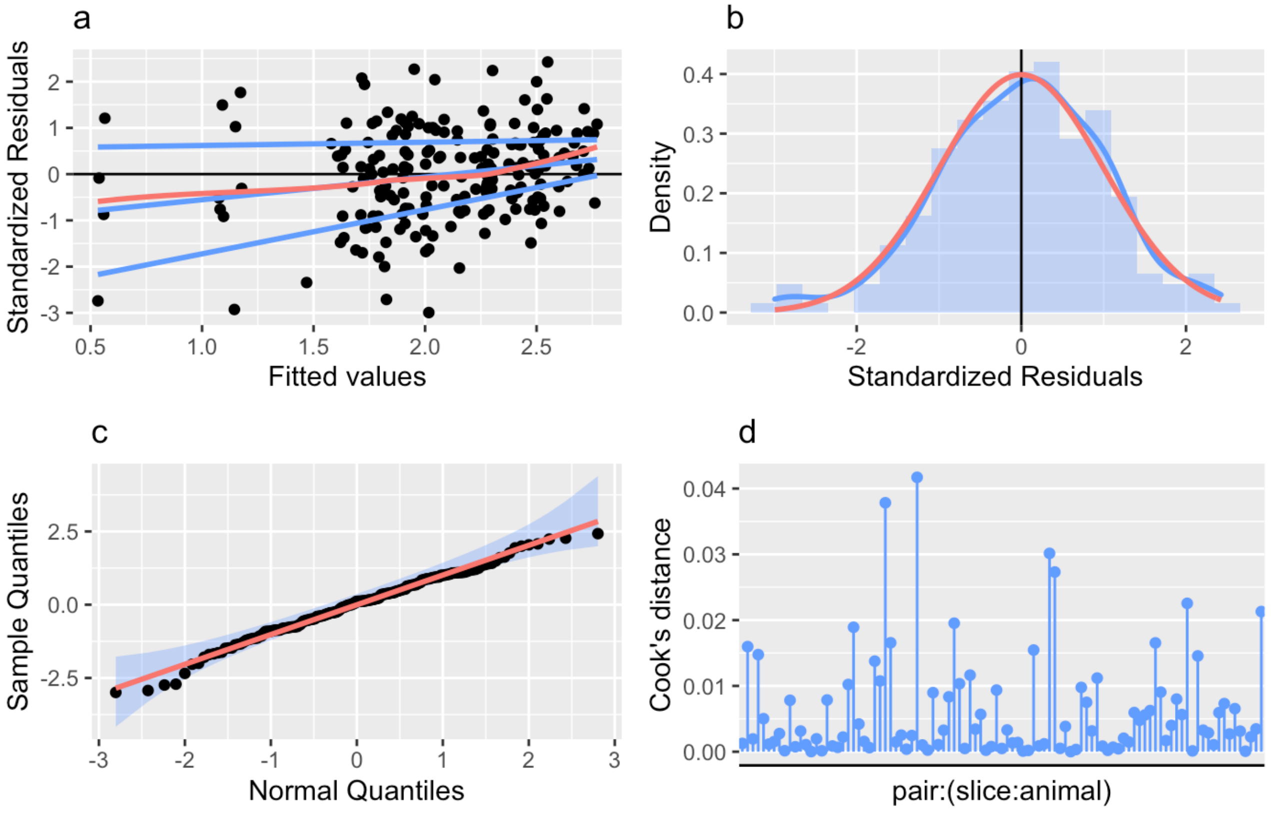

***Figure S1.9. NMDA-EPSC peak amplitudes in neurons expressing GluN2A knockout alleles.***

***Top***. Table summary of a LMM predicting NMDA-EPSC peak amplitude from the transfection of Cre-GFP in CA1 neurons of *grin2a$^{fl/fl}$* neurons. Fixed effects are summarised as an ANOVA table with the interaction term split into polynomial contrasts and followed up with posthoc pairwise comparisons. The *p*-value and Bayes factor for the interaction term omnibus test indicate a highly significant effect of loss-of-function (null) GluN2A alleles on the peak amplitude of NMDA-EPSCs. Of the variance not accounted for by the fixed effects, 27%, 39% and 44% was explained by variability between slices, animals and residual (unexplained) variance respectively. ***Bottom***. Model residuals appeared to be homoscedastic (**a**) and normally distributed (**b**, **c**), without outliers (**a, c**) or influential data points (**d**). A breakdown of samples sizes is reported in Table S4.

| **Response** | | | | | |
|---|---|---|---|---|---|
| NMDA-EPSC peak amplitude (peak) | | | | | |
| **Formula** | | | | | |
| $\log(\boldsymbol{peak}) \sim genotype * transfection + (1\vert animal/slice/pair)$ | | | | | |
| **Source (Fixed)** | ***F*** | ***df*** | ***$df_{res}$*** | ***p*** | ***$BF_{10}$*** |
| genotype | 0.83 | 2 | 10.07 | | |
| transfection | 25.46 | 1 | 96 | | |
| genotype:transfection | 8.61 | 2 | 96 | <.001 | 227 |
| Linear | 13 | 1 | 96 | <.001 | |
| Quadratic | 3.76 | 1 | 96 | .06 | |
| **Posthoc test** | ***t*** | | ***df*** | ***$p_{adj}$*** | |
| +/+ vs. +/- | -0.01 | | 96 | .99 | |
| +/+ vs. -/- | -3.61 | | 96 | .001 | |
| +/- vs. -/- | -3.47 | | 96 | .001 | |
| **Source (random)** | ***N*** | ***ICC*** | | | |
| pair:(slice:animal) | 99 | .00 | | | |
| slice:animal | 53 | .27 | | | |
| animal | 14 | .29 | | | |
| Residual | | .44 | | | |

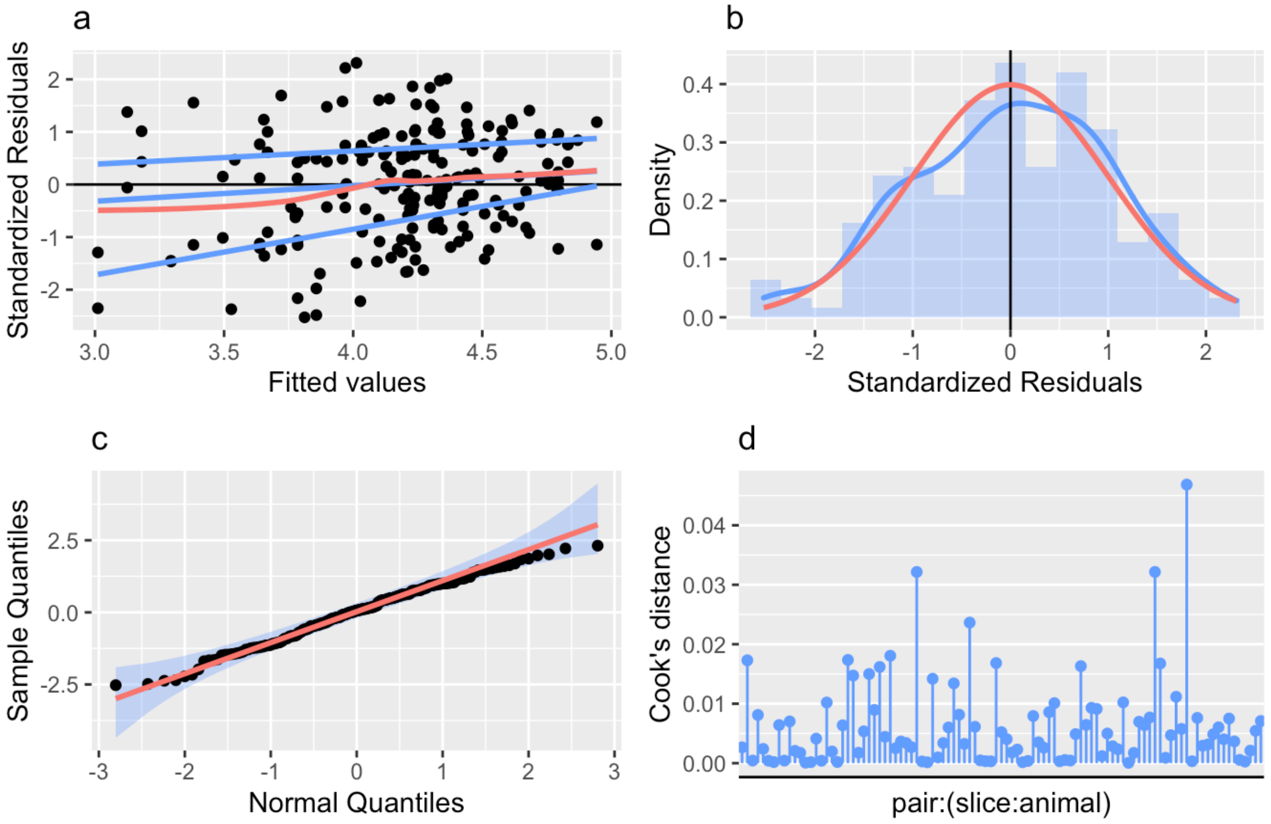

***Figure S1.10. NMDA-EPSC decays constant in neurons expressing GluN2A knockout alleles.***

***Top***. Table summary of a LMM predicting NMDA-EPSC decay time constant amplitude from the transfection of Cre-GFP in CA1 neurons of *grin2a*$^{fl/fl}$ neurons. Fixed effects are summarised as an ANOVA table with the interaction term split into polynomial contrasts and followed up with posthoc pairwise comparisons. The *p*-value and Bayes factor for the interaction term omnibus test indicate a highly significant effect of loss-of-function (null) GluN2A alleles on the decay time constant of NMDA-EPSCs. Of the variance not accounted for by the fixed effects, 14%, 26%, 25% and 35% was explained by variability between recording pairs, slices, animals and residual (unexplained) variance respectively. ***Bottom***. Model residuals appeared to be homoscedastic (**a**) and normally distributed (**b**, **c**), without outliers (**a, c**) or influential data points (**d**). A breakdown of samples sizes is reported in Table S4.

| **Response** | | | | | |
|---|---|---|---|---|---|
| NMDA-EPSC decay time constant (decay) | | | | | |
| **Formula** | | | | | |
| $\log(\boldsymbol{decay}) \sim genotype * transfection + (1 \vert animal/slice/pair)$ | | | | | |
| **Source (Fixed)** | ***F*** | ***df*** | ***df$_{res}$*** | ***p*** | ***BF$_{10}$*** |
| genotype | 4.91 | 2 | 9.90 | | |
| transfection | 156.8 | 1 | 96 | | |
| genotype:transfection | 76.23 | 2 | 96 | <.001 | 4.67E+19 |
| Linear | 140.1 | 1 | 96 | <.001 | |
| Quadratic | 9.9 | 1 | 96 | .002 | |
| **Posthoc test** | ***t*** | | ***df*** | ***p$_{adj}$*** | |
| +/+ vs. +/- | 2.84 | | 96 | .006 | |
| +/+ vs. -/- | 11.84 | | 96 | <.001 | |
| +/- vs. -/- | 8.52 | | 96 | <.001 | |
| **Source (random)** | ***N*** | **ICC** | | | |
| pair:(slice:animal) | 125 | .14 | | | |
| slice:animal | 78 | .26 | | | |
| animal | 27 | .25 | | | |
| Residual | | .35 | | | |

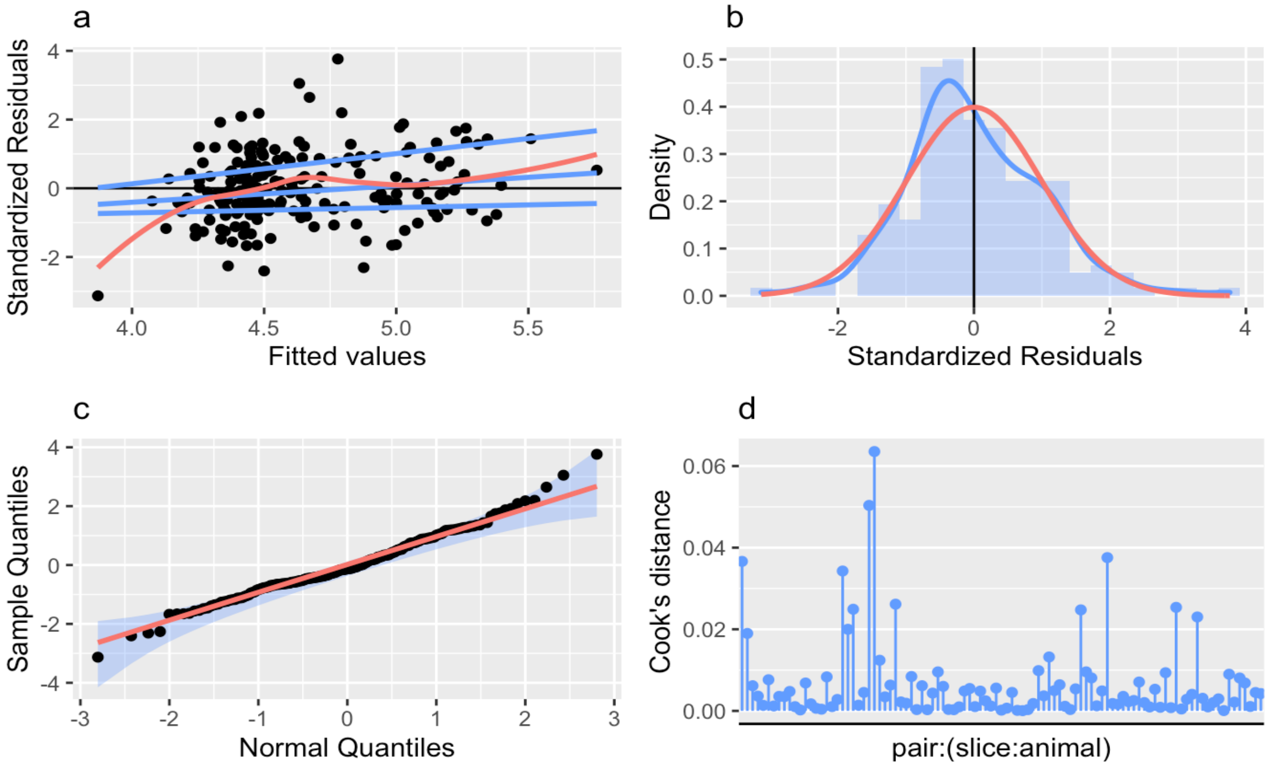

##### *Figure S1.11. NMDA-EPSC risetimes in neurons expressing GluN2A knockout alleles.*

***Top***. Table summary of a LMM predicting NMDA-EPSC 20-80% risetime from the transfection of Cre-GFP in CA1 neurons of *grin2a*$^{fl/fl}$ neurons. Fixed effects are summarised as an ANOVA table with the interaction term split into polynomial contrasts and followed up with posthoc pairwise comparisons. The *p*-value and Bayes factor for the interaction term omnibus test indicate a highly significant effect of loss-of-function (null) GluN2A alleles on the risetime of NMDA-EPSCs. Of the variance not accounted for by the fixed effects, 4%, 49%, 15% and 32% was explained by variability between recording pairs, slices, animals and residual (unexplained) variance respectively. ***Bottom***. Model residuals appeared to be homoscedastic (**a**) and normally distributed (**b**, **c**), without outliers (**a**, **c**) or influential data points (**d**). A breakdown of samples sizes is reported in Table S4.

| | | | | | |
|---|---|---|---|---|---|
| **Response** | | | | | |
| NMDA-EPSC 20-80% risetime (rise) | | | | | |
| **Formula** | | | | | |
| $\log(\boldsymbol{rise}) \sim genotype * transfection + (1 \mid animal/slice/pair)$ | | | | | |
| **Source (Fixed)** | ***F*** | ***df*** | ***df$_{res}$*** | ***p*** | ***BF$_{10}$*** |
| genotype | 1.5 | 2 | 9.69 | | |
| transfection | 83.79 | 1 | 96 | | |
| genotype:transfection | 44.21 | 2 | 96 | <.001 | 2.33E+13 |
| Linear | 81.48 | 1 | 96 | <.001 | |
| Quadratic | 5.55 | 1 | 96 | .021 | |
| **Posthoc test** | ***t*** | | ***df*** | ***p$_{adj}$*** | |
| +/+ vs. +/- | 1.07 | | 96 | .03 | |
| +/+ vs. -/- | 1.3 | | 96 | <.001 | |
| +/- vs. -/- | 1.22 | | 96 | <.001 | |
| **Source (random)** | ***N*** | **ICC** | | | |
| pair:(slice:animal) | 125 | .04 | | | |
| slice:animal | 78 | .49 | | | |
| animal | 27 | .15 | | | |
| Residual | | .32 | | | |

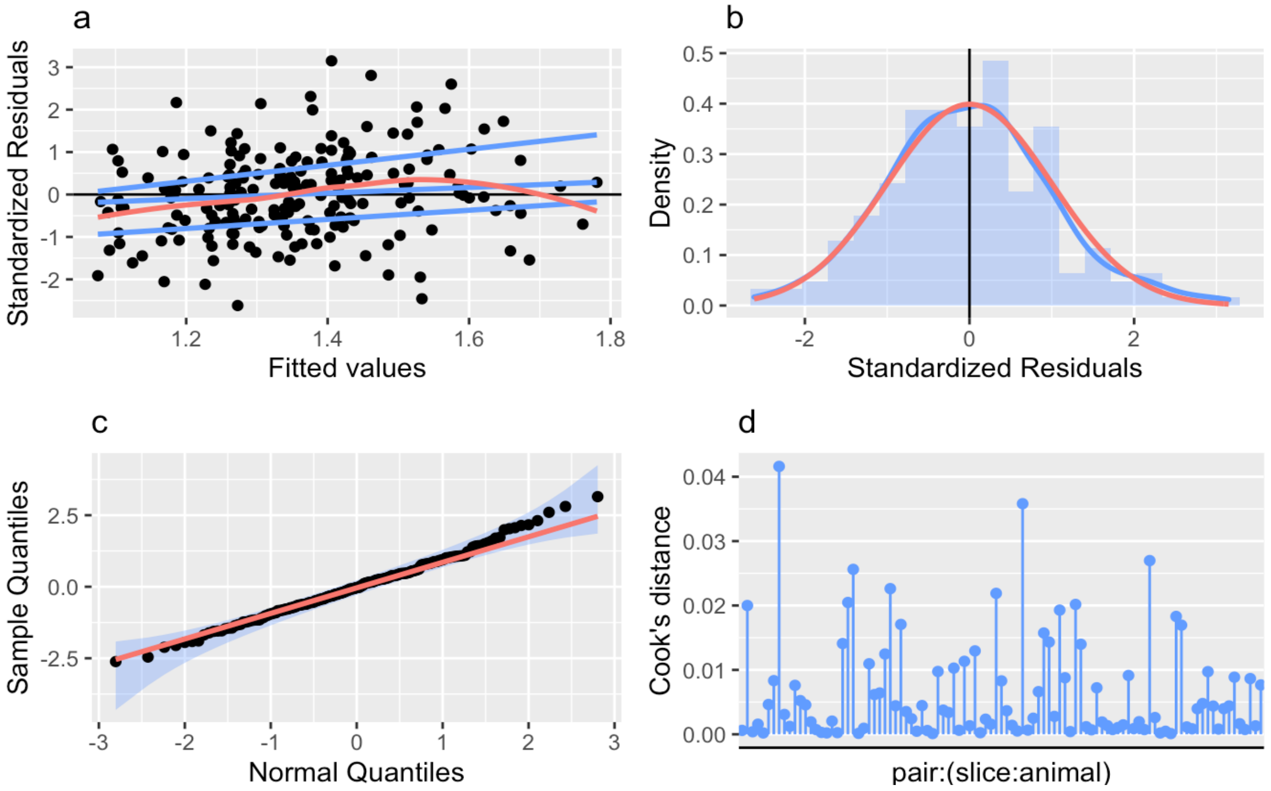

#### Figure S2: Likely toxicity from overexpressing the GluN2A L812M mutant

Example low magnification images and quantification of transfected cell density for GluN2A WT or L812M overexpressed in cultured hippocampal neurons. NMDA receptor blocker memantine (Mem) was effective at rescuing the density of neurons transfected with GluN2A L812M. The data points are plotted together with a bar graph representing the geometric mean and 95% confidence intervals.

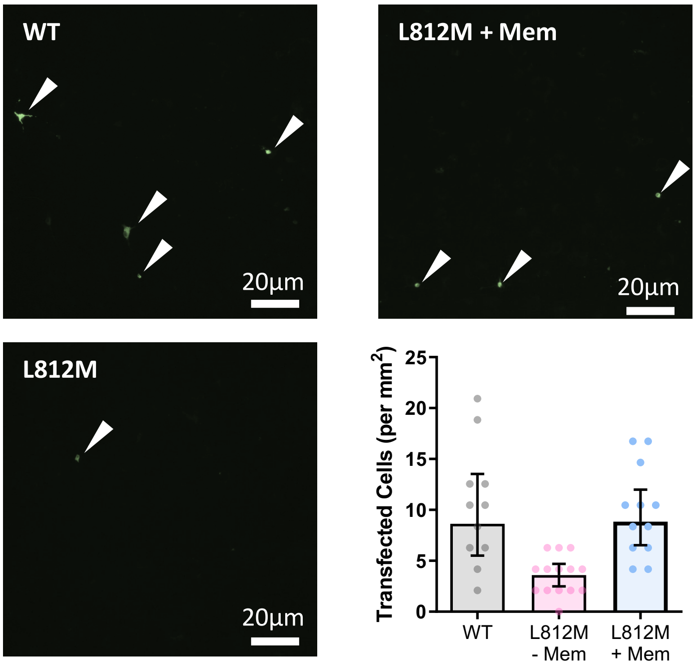

#### Figure S3: Effective rescue of native mouse GluN2A by human GluN2A

**a)** The experimental approach used to investigate the effect of missense mutations, by co-expressing Cre-GFP with human *GRIN2A* cDNA by single-cell electroporation around day 7 *in vitro*. The action of Cre recombinase results in the decline of native mouse GluN2A expression in Cre-transfected neurons of organotypic slices from *grin2a*$^{fl/fl}$ (as well as GluN2B in *grin2a*$^{fl/fl}$ *b*$^{fl/fl}$) mice slices. Meanwhile, native GluN2A is replaced by the co-expressed human GluN2A.

**b)** The transfected/untransfected ratios for NMDAR-EPSC peak amplitude and weighted decay time constants for varying amounts of co-transfected human *GRIN2A* cDNA. Arrow indicates the concentration that effectively rescued the peak amplitude and decay time constant of NMDAR-EPSCs. Error bars are 95% CI. The grey arrow indicates the concentration of *GRIN2A* cDNA used to rescue NMDA-EPSCs in experiments relating to Figures 2c, 2d, 4 and S3. The number of cell pairs recorded were 9, 14, 15, 15, 16 and 16 for 1.1, 3.3, 3.8, 4.4, 5.6 and 10.9 ng/µL respectively.

**c)** Example traces of NMDA-EPSCs measured at +20 mV to illustrate rescue of peak amplitude (*top*) and peak-scaled to illustrate rescue of the decay kinetics (*bottom*), of GluN2AKO cells by human GluN2A.

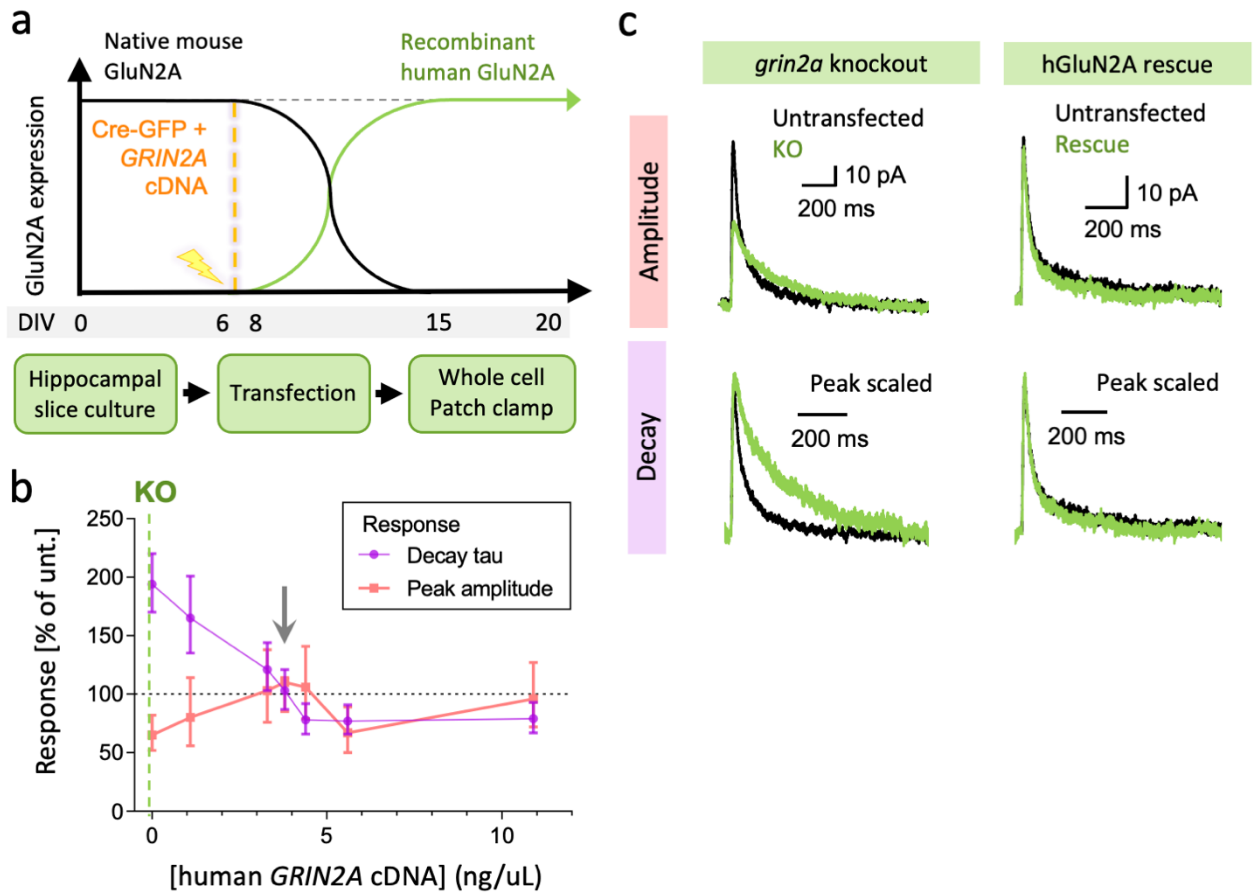

#### Figure S4: GluN2A mutations are not associated with large effects on AMPAR-EPSCs.

NMDA-EPSC$_{+20\ mV}$ peak amplitudes in *grin2a* $^{fl/fl}$ (untransfected) neurons and *grin2a* $^{-/-}$ neurons rescued with human GluN2A WT, LOF (C436R, T531M or R518H) or GOF (K669N or L812M) mutants (transfected). **ai)** Data points of measurements made in individual neurons. Matched data points, for simultaneously recorded untransfected and transfected neurons, are connected by a line. **aii)** Response ratios (transfected/untransfected) are expressed as a percentage and plotted for each pair of transfected-untransfected neurons. Crossbars in **i)** and **ii)** show the estimated marginal means with 95% confidence intervals backtransformed from the linear mixed models (Figure S11). Hypothesis tests are orthogonal contrasts based on *a priori* clustering of the mutations in Figure 1e. Standardized effect sizes (*r*) for comparisons of each mutant with WT for response ratios of peak amplitude in **aii** were .09, .07, .05, -.08 and -.08 (*N* = 125).

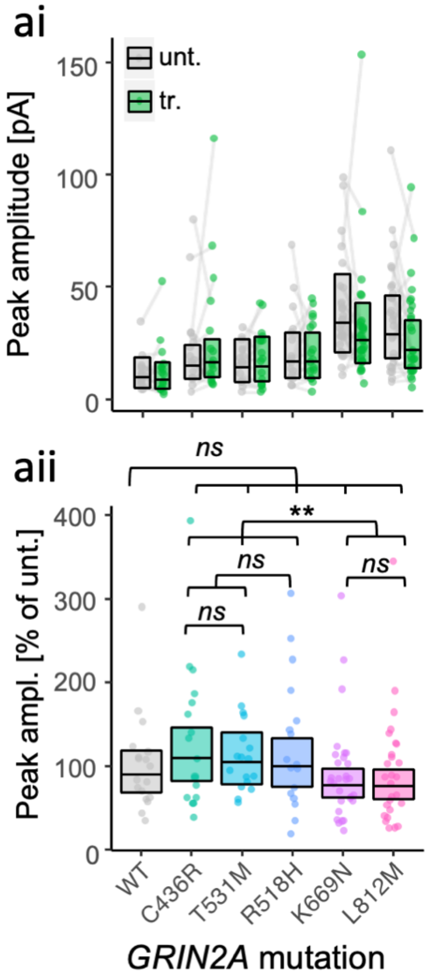

#### Figure S5: Structural models of the wildtype and mutated amino acids for LOF mutations

C436R disrupts a disulphide bridge. T531M modifies interactions around the LBD hinge between the upper and lower lobes of the clamshell. R518H disrupts key coulombic and hydrogen bonding interactions between the glutamate (Glu) ligand and the LBD.

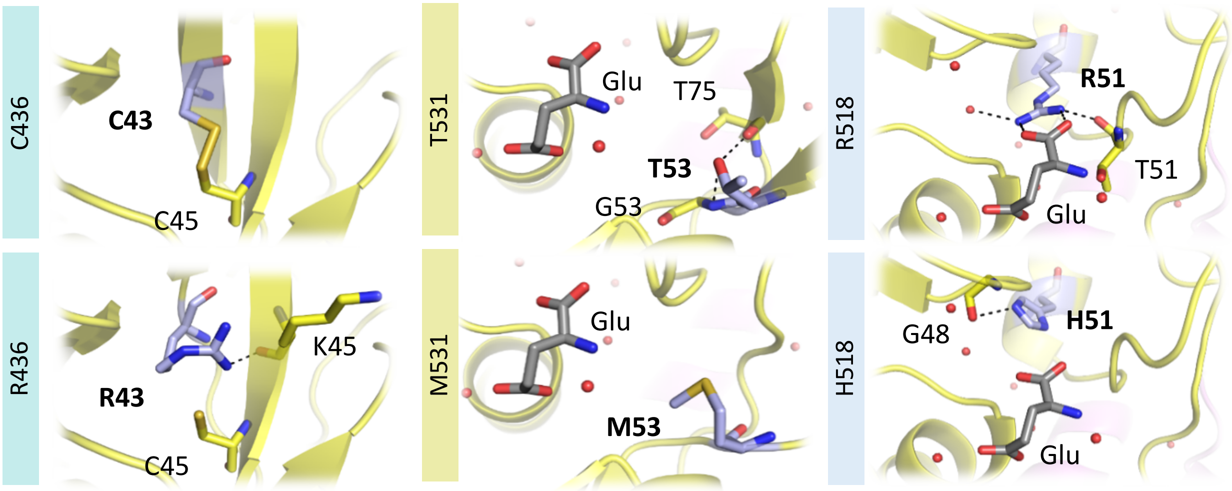

#### Table S1. Matrix of orthogonal contrasts based on *a priori* clustering of mutations.

Matrix of orthogonal contrasts used in the analysis of data in Figures 2ciii, 3bii, 4aiii, 4biii and 4ciii. Conditional table formatting is used to colour code the mutants involved in each contrast and illustrate their weighting.

| **Contrast** | **A** | **B** | **C** | **D** | **E** |
|---|---|---|---|---|---|
| ***GRIN2A*** | **WT *vs* Mutants** | **LOF *vs* GOF** | **LOF2 *vs* LOF1** | **LOF2** | **GOF1** |
| **WT** | -0.83 | 0 | 0 | 0 | 0 |
| **C436R** | 0.17 | -0.4 | -0.33 | -0.5 | 0 |
| **T531M** | 0.17 | -0.4 | -0.33 | 0.5 | 0 |
| **R518H** | 0.17 | -0.4 | 0.67 | 0 | 0 |
| **K669N** | 0.17 | 0.6 | 0 | 0 | -0.5 |
| **L812M** | 0.17 | 0.6 | 0 | 0 | 0.5 |

#### Table S2: Clinical features of patients reported previously with *GRIN2A* mutations used in this study.

| *GRIN2A* mutation | Epilepsy Syndrome | Seizures | EEG | Speech | DD | ID | MRI | Ref |
|---|---|---|---|---|---|---|---|---|
| **C436R** | ABPE | Nocturnal GTCS, atonic seizures | CTS | Mild speech delay | Normal | Normal | ND | [1] |
| **T531M** | ECSWS, IEAD | Focal Dyscognitive seizure, Tonic Clonic Seizures | Intermittent discharges in clusters - LPH, HVBD (awake), Bilateral CTS (drowsiness & sleep), Multifocal discharges - mid parietal, independent bilateral bioccipital regions. Very frequent in awake state and continuous in sleep | Speech / Language Difficulties | Mild, Global | Mild, Learning difficulties | Normal | [2] |
| **R518H** | CSWSS, Landau-Kleffner Syndrome, atypical RE | Partial seizures | ESES, CTS | verbal dyspraxia | Normal | ND | Normal | [3] |
| **K669N** | CSWSS | Partial seizures | ESES | Language regression | Profound, Global | ND | Normal | [3] |
| **L812M** | Intractable infantile-onset epilepsy | Daily, Generalised seizures | Diffusely slow, disorganized background, irregularly generalized/lateralized spike and slow wave discharges, slow activity (3.5–4.5 Hz, 1–3 Hz delta and disorganized activity), intermittent irregular high amplitude 2.5 Hz spike-wave discharges (frontal, greater on right hemisphere). | ND | Profound, Global | Profound | Diffuse cerebral parenchymal volume loss, thin corpus callosum | [4,5] |

ABPE - Atypical benign partial epilepsy, CSWSS - continuous spike-wave of slow sleep, CTS – centrotemporal spikes, DD – developmental delay, ECSWS - epileptic encephalopathy with continuous spike-and-wave during sleep syndrome, EEG – electroencephalogram, ESES - Electrical status epilepticus during slow-wave sleep, GTCS - generalized tonic-clonic seizure, HVBD - High voltage bilateral discharges, ID – intellectual disability, IEAD - intermediate epilepsy-aphasia disorder, LPH – Left posterior hemisphere, MRI – magnetic resonance imaging, ND – Not determined , RE – Rolandic Epilepsy

#### Table S3: Primers used to generate mutant GluN2A constructs from WT pCI-Neo *GRIN2A*

| ***GRIN2A*** **MUTATION** | **Forward primer (5' to 3')** | **Reverse primer (5' to 3')** |
|---|---|---|
| **C436R** | AGGAACACCGTGCCACGTCGGAAGTTCGTCA | TGACGAACTTCCGACGTGGCACGGTGTTCCT |
| **R518H** | CCATCAATGAGGAACATTCTGAAGTGGTGGA | TCCACCACTTCAGAATGTTCCTCATTGATGG |
| **T531M** | TGCCCTTTGTGGAAATGGGAATCAGTGTCAT | ATGACACTGATTCCCATTTCCACAAAGGGCA |
| **K669N** | CCTCAGTGACAAAAATTTTCAGAGACCTCAT | ATGAGGTCTCTGAAAATTTTTGTCACTGAGG |
| **L812M** | GTGATGAGCAGCCAGATGGACATTGACAACA | TGTTGTCAATGTCCATCTGGCTGCTCATCAC |

#### Table S4. Experiment sample sizes

Sample sizes for experiments detailed in Figures S1.1-1.11 broken down into the number of recording (cell) pairs, number of slices and number of animals.

| Experiment relating to Figure 2b | | | |
|---|---|---|---|
| **Condition** | **pairs** | **slices** | **animals** |
| DKO | 19 | 11 | 3 |
| DKO DQP | 15 | 9 | 3 |

| Experiment relating to Figure 2c, d | | | |
|---|---|---|---|
| Mutation | pairs | slices | animals |
| WT | 14 | 8 | 3 |
| C436R | 19 | 12 | 3 |
| T531M | 9 | 5 | 2 |
| R518H | 13 | 8 | 2 |
| K669N | 21 | 10 | 3 |
| L812M | 17 | 11 | 3 |
| WT(2B) | 19 | 13 | 3 |

| Experiment relating to Figure 3b | | |
|---|---|---|
| Mutation | roi | expt |
| WT | 57 | 6 |
| C436R | 20 | 3 |
| T531M | 41 | 3 |
| R518H | 33 | 3 |
| K669N | 22 | 3 |
| L812M | 18 | 3 |
| NONE | 81 | 7 |

| Experiment relating to Figure 4 | | | |
|---|---|---|---|
| Mutation | pairs | slices | animals |
| WT | 18 | 12 | 3 |
| C436R | 17 | 12 | 6 |
| T531M | 17 | 9 | 3 |
| R518H | 17 | 11 | 4 |
| K669N | 29 | 15 | 5 |
| L812M | 27 | 19 | 6 |

| Experiment relating to Figure 5 | | | |
|---|---|---|---|
| Genotype | pairs | slices | animals |
| WT | 33 | 19 | 4 |
| HET | 29 | 15 | 6 |
| HOM | 37 | 19 | 4 |

#### Supplemental Methods

##### *Structural modelling*

Structural modelling was performed on GluN2A in a pseudo full-length human model of a triheteromeric NMDA receptor. The coordinates of the triheteromer structure of *Xenopus laevis* GluN1/2A/2B (4.5Å, 5uow) were downloaded from the Orientations of Proteins in Membranes (OPM) database[6]. GalaxyFill was used to fill in missing main chain and side chain atoms and loops of each protomer[7]. The protomers then the reassembled and steric clashes removed by brief energy minimization using steepest descent. This structure had a root mean squared deviation of CA atoms (CA-RMSD) of 0.08 Å (from 5uow) and was used as scaffold for modelling the ligand-bound human triheteromeric receptor in Modeller (v 9.19)[8] using multiple high-resolution templates: 4nf8, 5fxh, 5fxi, 5i57, 5tpw, 5tpz and 5un1. The *automodel* function was used to create 400 models using 1000 iterations of the variable target function method (VTFM, *autosched.slow*) followed by Simulated Annealing Molecular Dynamics (SA-MD) refinement (*refine.very_slow*). Each model was assessed using three statistically optimized atomic potentials (SOAP, soap_protein_od, soap_pp and soap_peptide) to assess sidechain packing, subunit interaction surfaces and amino acid ligands[9]. Following transformation of each potential to Z-scores, models were accepted whose scores were more negative than -1 standard deviation. Of the accepted models, the one with the lowest combined potential was taken forward for refinement by restrained molecular dynamics in CHARMM[10] using the CHARMM36m forcefield[11]. In brief, alternative protonation states for Glu, Asp and Lsn (at neutral pH) were identified using PROPKA3.1[12] and patched along with disulphide bridges and C-terminal *N*-methylamide. Hydrogens were added using H-BUILD and the structure was embedded in a heterogeneous dielectric generalized Born (HDGB) implicit membrane (v3)[13] with infinite cut-offs for non-bonded interactions. Solvent accessible residues within the channel were excluded from the low dielectric using the EXCLGB parameter[14]. Within the implicit solvent and membrane environment, the structure was subjected to a series of restrained energy minimization and molecular dynamics using a protocol closely resembling the Local Protein Structure Refinement via Molecular Dynamics Simulations (locPREFMD)[15]. Molecular dynamics simulations were performed using the velocity Verlet integrator with a 1 fs time step and SHAKE constraints applied on bonds involving hydrogen.

*In silico* point mutations were generated in the NMDA receptor model using Modeller 9.19 by a combination of conjugate gradient minimization and molecular dynamics simulated annealing[16] (https://salilab.org/modeller/wiki/Mutate model). The procedure was repeated with different random seeds to generate 100 different unique models and the model with the lowest energy was selected.

All modelling and refinement steps were accomplished on a Dell Precision T7910 workstation with dual Intel Xeon processors (E5-2687W @ 3.10 GHz, 10 cores each), Structures were visualized (and figures created) in Pymol (The PyMOL Molecular Graphics System, Version 1.7.7.6 Schrödinger, LLC.).
